# Ranging patterns of the rainforest-adapted lion-tailed macaque *Macaca silenus* in a human-dominated landscape in the Anamalai hills of the Western Ghats, India

**DOI:** 10.1101/2022.08.04.502767

**Authors:** Ashni Kumar Dhawale, Anindya Sinha

**Affiliations:** Animal Behaviour and Cognition Programme, National Institute of Advanced Studies, Bangalore, India; University of Trans-Disciplinary Health Sciences and Technology, Bangalore, India; Indian Institute of Science Education and Research, Kolkata, India; Centre for Wildlife Studies, Bangalore, India

## Abstract

The ranging patterns of five lion-tailed macaque *Macaca silenus* troops, forming the Puthuthottam sub-population, were studied over a three year period to determine road/habitation visitation rate, home ranges and habitat preference. Each troop visited the road or human habitation at varying rates, with the largest troop visiting most frequently. Home ranges sizes were observed to be highly reduced when compared to wild populations, and also greatly varied across troops, with relatively low overlap given the macaque density in the available area. All five macaque troops showed a preference for human-modified habitats such as roads and human settlements where anthropogenic food was easily available. Our study shows an increasing dependence amongst members of the Puthuthottam troops on anthropogenic foods, which has led to many threats faced by individuals including fatal collisions with vehicular traffic and electrocutions.

## Introduction

Animal movements in a landscape are largely determined by the availability and distribution of food, predation risk, intra- and inter-species competition, reproductive investments and behavioural adaptations (Clutton-Brock, 1975; Pontzer & Kamilar, 2009; South, 1999; Wiens et al., 1993), all of which are heavily influenced by human disturbance. Globally, natural habitats are being razed for agricultural purposes, having already resulted in the loss of up to 50% of forest land (Defries et al., 2004). The resultant fragmentation of natural habitats into isolated patches is known to drastically affect the spatial movement of animals, often restricting them to certain areas beyond which the habitat becomes impermeable (Andren, 1994; Bladon et al., 2002; Mbora et al., 2009). Patch resource quality and heterogeneity directly influences animal home range (Levins, 1968; Rolstad, 1999), which is typically defined as the total area used by an individual or group (Jay, 1965). Additionally, fragmentation indirectly shapes ranging behaviour through the introduction of unnatural features into the landscape, such as linear intrusions and barriers (Jakes et al., 2018). Ranging behaviour, thus, becomes a useful tool to capture the interactions of animals with their changing environment, especially insofar as human activity is concerned.

Humans developmental activities directly and indirectly impact the ranging behaviour of animals. For example, the construction in windfarms in Scotland caused resident Golden eagles *Aquila crysaetos* to change their ranging in order to avoid the manmade structures (Walker et al., 2005). The red fox *Vulpes vulpes* selectively used human-dominated areas in Central Italy, based on the tolerance exhibited by people towards them (Lucherini et al., 1995). Another species from Central Italy, the Least weasel *Mustela nivalis* showed a strong preference for remnant natural habitats, such as hedges, in a predominantly agricultural landscape (Magrini et al., 2009). A species of stone curlew *Burhinus oedicnemus* in southern England preferentially chose breeding grounds that were greater than three kilometres from a major road (R. E. Green et al., 2000). Home range expansion was observed to be multi-fold in some populations of South Andean deer *Hippocamelus bisulcus* in response to hunting and other such human disturbances in Chilean Patagonia (Gill et al., 2008). Closer to home, in the Western Ghats of India, movement patterns of many large mammals including the Asian elephant *Elephas maximus*, spotted deer *Axis axis* and tiger *Panthera tigris* are radically affected by linear intrusions such as pipelines, railway tracks, electric wires and fences (Menon et al., 2013; Nayak et al., 2020).

Of the many mammals impacted by human activity, primates perhaps have the longest history of interactions with humans and human habitations. For example, the Bonnet macaque *Macaca radiata*, endemic to peninsular India, has featured in literature from 2000 years ago, being described as a regular fixture in the town’s commons (see Sinha, 2001 for source). With a population of 2 billion people in primate range countries, as of 2005 (Estrada et al., 2012), it is hardly surprising that primates across the world are increasingly encountering humans and their infrastructure. In fact, many species the world over, are able to persist in agroecosystems, or habitats dominated by crops but having some remnant natural vegetation (Estrada et al., 2006, 2012). These trends, however, are usually observed in primate species that show a high propensity for adapting to human-dominated landscapes, such as habitat generalists or species that are non-reliant on dense canopy for movement. Even those that find their way through a human-modified habitat matrix, and are able to exploit new food sources or find shelter (Adhikari et al., 2018; Estrada et al., 2012; Ganguly & Chauhan, 2018; Nijman, 2021), face numerous caveats including intra-species and human-primate conflict (Defries et al., 2004; Jaman & Huffman, 2013; Radhakrishna & Sinha, 2011; Ram et al., 2003; Riley, 2007; Sinha et al., 2005; Tracie, 2011; Warren et al., 2011), fatal encounters with vehicles, increased parasite load (Hussain et al., 2013; Mbora et al., 2009) and hunting pressures (Gill et al., 2008; Richard-Hansen, 2000).

These caveats are especially pronounced in those primate species that display a further specialisation in their ecology or behaviour. For example, the highly arboreal proboscis monkey *Nasalis larvatus* completely avoided clear-felled habitats surrounding human habitation (Salter et al., 1985) and abandoned roosting sites along riversides where tourism-associated infrastructure was established (Marsh & Chapman, 2013). The Yunnan snub-nosed monkey, inhabiting the highest elevation of any non-human primate species, displayed greatly varied daily movements in response to severe human disturbance, which were further exacerbated by the seasonality of natural food resources (Li et al., 2020). The habitat-specialist diademed sifaka *Propithecus diadema* showed a drastically reduced home range size and daily path length in fragmented habitats (Irwin, 2008). A similar trend was observed in frugivorous primates, such as the moustached guenon *Cercopithecus cephus* and hoolock gibbon *Hylobates hoolock* and *Hylobates agilis* (Yanuar & Chivers, 2010), wherein home range size in fragmented habitats is drastically reduced. This pattern could perhaps be explained by the surrounding human-dominated matrix creating a “hard edge”, restricting the species entirely within the fragment, as is the case with the highland mangabey *Rungwecebus kipunji* in Tanzania (Bracebridge et al., 2013). It is also noteworthy that folivorous species that have inherently small home ranges tend to fare better in fragmented habitats, as they are able to maximise resources within restricted areas (Yanuar & Chivers, 2010).

The lion-tailed macaque *Macaca silenus*, while belonging to the highly adaptable genus of macaques, has been categorised as an arboreal, primarily frugivorous, habitat-specialist species, dependent on the wet evergreen native vegetation type (Kumar, 2013). This is a species endemic to the Western Ghats, existing today in 49 subpopulations, in only eight key locations, including the Anamalai Hills (Kumara & Singh, 2003; Kurup & Kumar, 1993; Molur et al., 2003). Since the late 1800s, logging of the native vegetation for the expansion of commercially grown tea and coffee plantations on the Valparai plateau in the Anamalai hills has resulted in forest fragmentation and the isolation of lion-tailed macaque troops now scattered within these remaining pockets of rainforest (Jeganathan et al., 2018; Singh et al., 2002). Despite the degraded nature of these remaining habitats, the Anamalai hills, being contiguous with Parambikulam Tiger Reserve and Neliampathy in the North, and the Chalakudi hills in the south, has been identified as a crucial landscape for the conservation of the lion-tailed macaque (Singh et al., 2002).

The Valparai plateau is a matrix of tea and coffee plantations interspersed with 45 rainforest fragments ranging in size from <10ha to >100ha (Mudappa & Shankar Raman, 2007; Umapathy & Kumar, 2000). In the surrounding shola forest of Varagaliyar, lion-tailed macaque groups are reported to maintain a home range of 131ha, while covering 10.75ha and moving between 0.75km to 2.5km on a daily basis (Kurup & Kumar, 1993). In contrast, this study focuses on one of the larger forest fragments in the Valparai plateau, measuring 92ha. The Puthuthottam forest fragment, neighbouring the town of Valparai, and surrounded on all other sides by tea-plantations, contains ∼190 lion-tailed macaque individuals divided into five troops. All of the five troops present in the Puthuthottam forest fragment visit human habitations, either labour lines within the fragment or the neighbouring town of Valparai (Dhawale Pers. Obs.).

Troops in this population already exhibit adaptations to these anthropogenic habitats, significantly reducing time spent foraging while increasing time spent resting, and display altered social dynamics under regimes of potentially perceived competition in the presence of human-use foods (Dhawale et al., 2020). Like many other macaque species (Greenwood, 1980), male lion-tailed macaques typically disperse from the natal troop at sexual maturity. This dispersal pattern is thought to reduce inbreeding in species (Moore, 1992), thus playing a crucial role in their long-term survival. In fragmented landscapes like the Valparai plateau, however, male migration in lion-tailed macaques is severely impeded (Singh et al., 2002). As a result, males tend to stay back in the natal troops, which has led to an unusual multimale/ multifemale social organisation in the troops present in Puthuthottam (Dhawale, pers. obs.). Given these relatively recent shifts in the species’ ecology and behaviour in this particular population, we sought to examine the ranging behaviour, and habitat use and preference of the five Puthuthottam troops, as they traversed over a human-dominated habitat matrix.

## Objective and Questions

Examining movement, habitat use and competition across the multiple lion-tailed macaque troops residing in Puthuthottam forest fragment through ranging behaviour.

1. How does the home range differ between troops and across field seasons?
2. How much overlap is observed across troop core- and outer-home ranges?
3. What is the degree of movement for each troop per day over the study period?
4. What pattern of habitat use is observed by the Puthuthottam population over the study period?

## Methods

### Field Methods

GPS locations were taken at the centre of two pre-determined marker adult females of the troop at 15 min intervals during the simultaneous and systematic following of all troops present in the Puthuthottam forest fragment, as they ranged over both natural and anthropogenic habitats. Data was collected for 8±1 months (over 14±1.7 days/ month) for each season (September to May), on each of the five troops present in Puthuthottam from 2018 to 2021. A total of ∼5000 location data points were collected with 250±100 data points per month.

#### Habitation Visitation Rate

The Puthuthottam highway and human settlements were monitored continuously for 12 months between October 2018-October 2019, with a GPS location recorded at every encounter with any troop present in Puthuthottam. If the troop continued to remain by the human habitation, the GPS record was repeated at 15 min intervals. These locations were later mapped to describe the patterns of road visitation by the lion-tailed macaques in the Puthuthottam population.

#### Home range estimation, overlap and habitat use

GPS locations over the field seasons and total study period were mapped using GIS software to calculate distances travelled, directions of movement and rates of ranging across and outside the study area for each troop. Such an analysis is essential to map the new-found home range of these macaque troops, particularly in so far as they overlap with human habitations, orchards and roads, potential areas for escalated human-primate conflict.

### Analysis

#### Habitation Visitation Rate

A habitation visitation rate was calculated as the proportion of days over the monitoring period during which lion-tailed macaques were encountered near roads or human settlements (adapted from Singh 2001).

#### Home range estimation and overlap

A kernel density estimation (Laver & Kelly, 2008) allowed us to determine outer home range (95% use area) and core use area (50% use area), using an optimal bandwidth selection method to delineate kernels from Fotheringham et al., 2000. KDE calculations and visualisation were completed in QGIS (QGis, 2011 version 2.18.3) using the Heatmap plugin. Additionally we visually present overlap of home range across all troops in Puthuthottam to describe prevailing inter-troop competition.

#### Degree of movement

Daily paths were calculated for each troop over each field season in QGIS (QGis, 2011 version 2.18.3) and their lengths presented as average per troop per field season.

#### Habitat Use and Preference

To describe habitat use and preference, troop locations were sampled such that a single unique location was considered per day over the entire study period, and compared to a randomly generated set of points of comparable sample size using a non-parametric test (Wilcoxon Test) in R, revised version 3.2.4. The random points were weighted as density dependent, based on the available area of any given habitat type using QGIS (QGis, 2011 version 2.18.3). Additionally, the study area was rasterized such that each raster pixel (50mx50m) contained a corresponding ‘Habitat Type’ value and the frequency of each habitat type (available area) was compared to the sampled troop locations to provide a visual comparison of availability versus use.

All graphs were created in R, revised version 3.2.4. The habitat types considered are as follows:

##### Forest Edge

A 50-m-wide belt around the edge of the Puthuthottam forest fragment, containing native and non-native tree species and bordered on one side by a national highway. We chose to demarcate the boundary at 50m from the edge as we observed that the troop spread at any given time was ≤ 50m. This habitat contained Natural food sources, and occasionally Human-use foods, either dropped along the roadside or in the form of handouts provided by tourists.

##### Forest Interior

An area of forest contained by the Forest Edge, described above, consisting of native and non-native tree species, all of which constituted Natural food sources.

##### Open Forest Patch

A relatively open space, largely without canopy cover, present within the Puthuthottam forest and recently planted with coffee saplings. It included only Natural food sources.

##### Human Settlement

Six separate human habitations were present within and surrounding the Puthuthottam forest fragment, including two high-density towns, three labour lines housing plantation workers, and a hospital building. These areas were considered as Human Settlement habitat type which was characterised by the presence of both Natural and Human-origin food resources.

##### Puthuthottam Road

The section of the Puthuthottam Highway beginning from the Human Settlements to the north of the forest fragment and ending at the southern end of the forest fragment.

## Results

### Habitation Visitation Rate

Each of the five troops visited habitation at varied rates (Figure 1), with the BT troop and RT troop visiting habitation most frequently. The overall habitation visitation rate was calculated to be 0.57 times a day.

**Fig 1.**
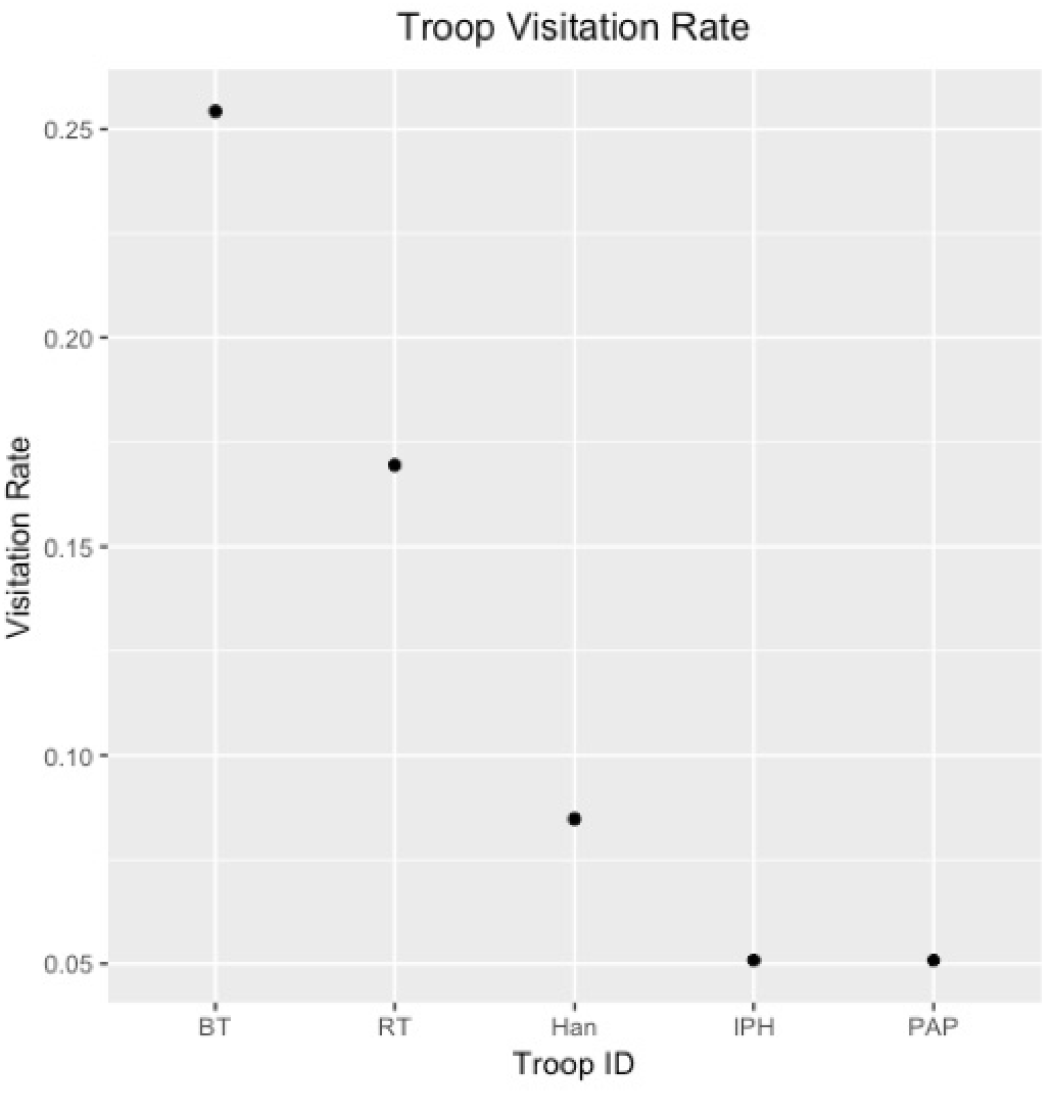
Habitat visitation rate of the five Puthuthottam troops during October 2018-October 2019

### Outer- and core-home range

Figure 2 depicts the home ranges for each field season (Sept 2018-May 2019; Aug 2019-Apr 2020; Oct 2020-Mar 2021) of all troops present in Puthuthottam. Three of the troops were present for only a few of the field seasons due to fission-fusion and reforming of certain troops. Home ranges varied both across field seasons and troops. The home range of the biggest troop, BT, seemed to expand over the three field seasons while the RT home range seemed to become concentrated in certain areas. NTT, a troop which formed when two smaller troops joined together, seemed to show the most varied home range over field seasons. Figure 3 depicts the overall home range of each troop measured over the entire study period. Three of the troops, namely BT, PAP and HAN ranged primarily over the southern part of the forest fragment, while the other two troops, NTT and RT, were mostly observed in the northern part of the fragment. Table 1 contains the overall home range sizes of each troop. The largest troop, BT, also had the largest home range, however, the smallest troop, HAN, did not have the smallest home range. Figure 4 shows the correlation between total home range area and total troop size. There is a slight correlation between home range and troop size (Spearman rank correlation, R= 0.6, p=0.35), and home range and number of adult males per troop (data not shown), however, these were not statistically significant.

**Table 1.**
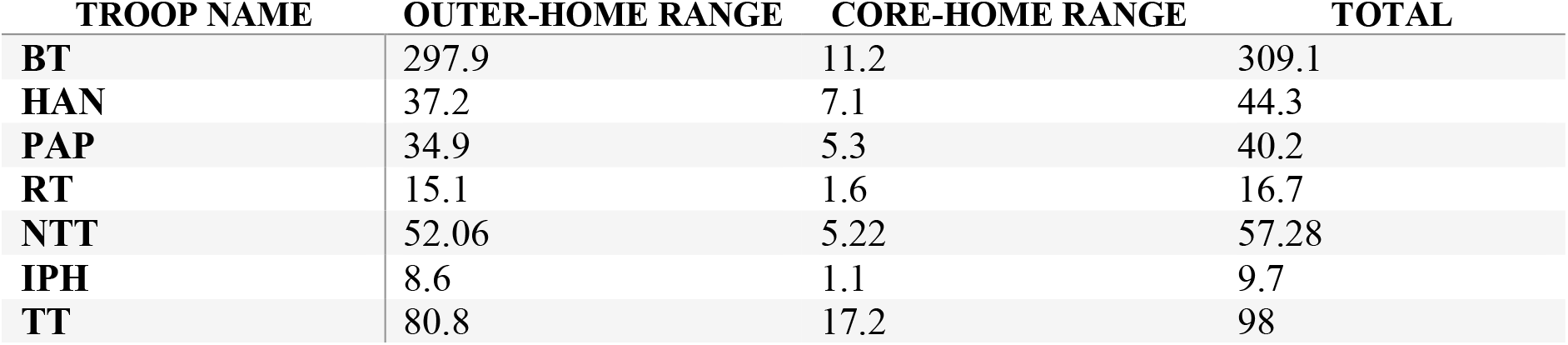
Overall home range sizes (ha) of all troops in Puthuthottam 2018-2021

| TROOP NAME | OUTER-HOME RANGE | CORE-HOME RANGE | TOTAL |
| --- | --- | --- | --- |
| BT | 297.9 | 11.2 | 309.1 |
| HAN | 37.2 | 7.1 | 44.3 |
| PAP | 34.9 | 5.3 | 40.2 |
| RT | 15.1 | 1.6 | 16.7 |
| NTT | 52.06 | 5.22 | 57.28 |
| IPH | 8.6 | 1.1 | 9.7 |
| TT | 80.8 | 17.2 | 98 |

**Fig 2.**
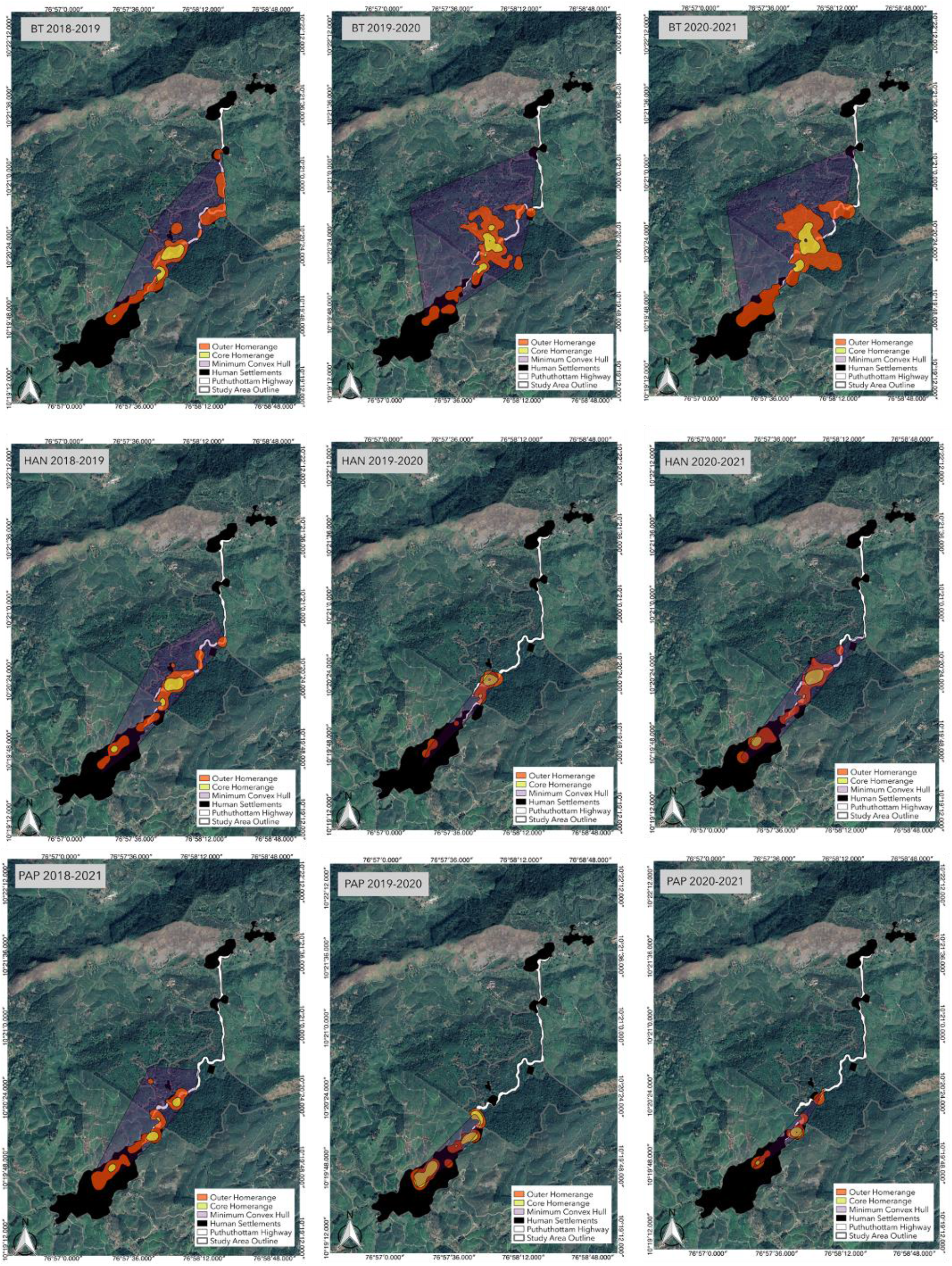
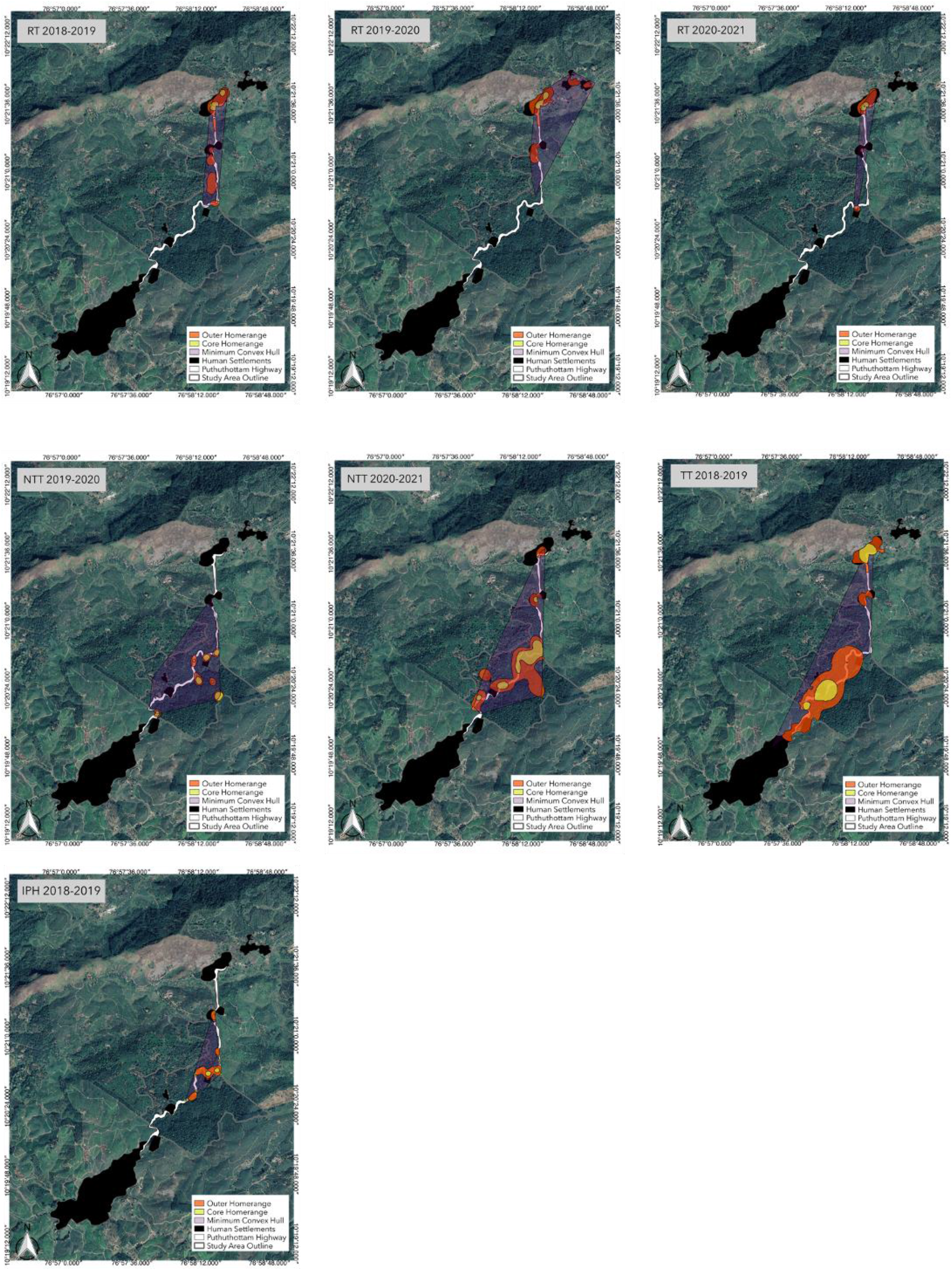
Field season-wise home range for each of the five troops present in Puthuthottam

**Fig 3.**
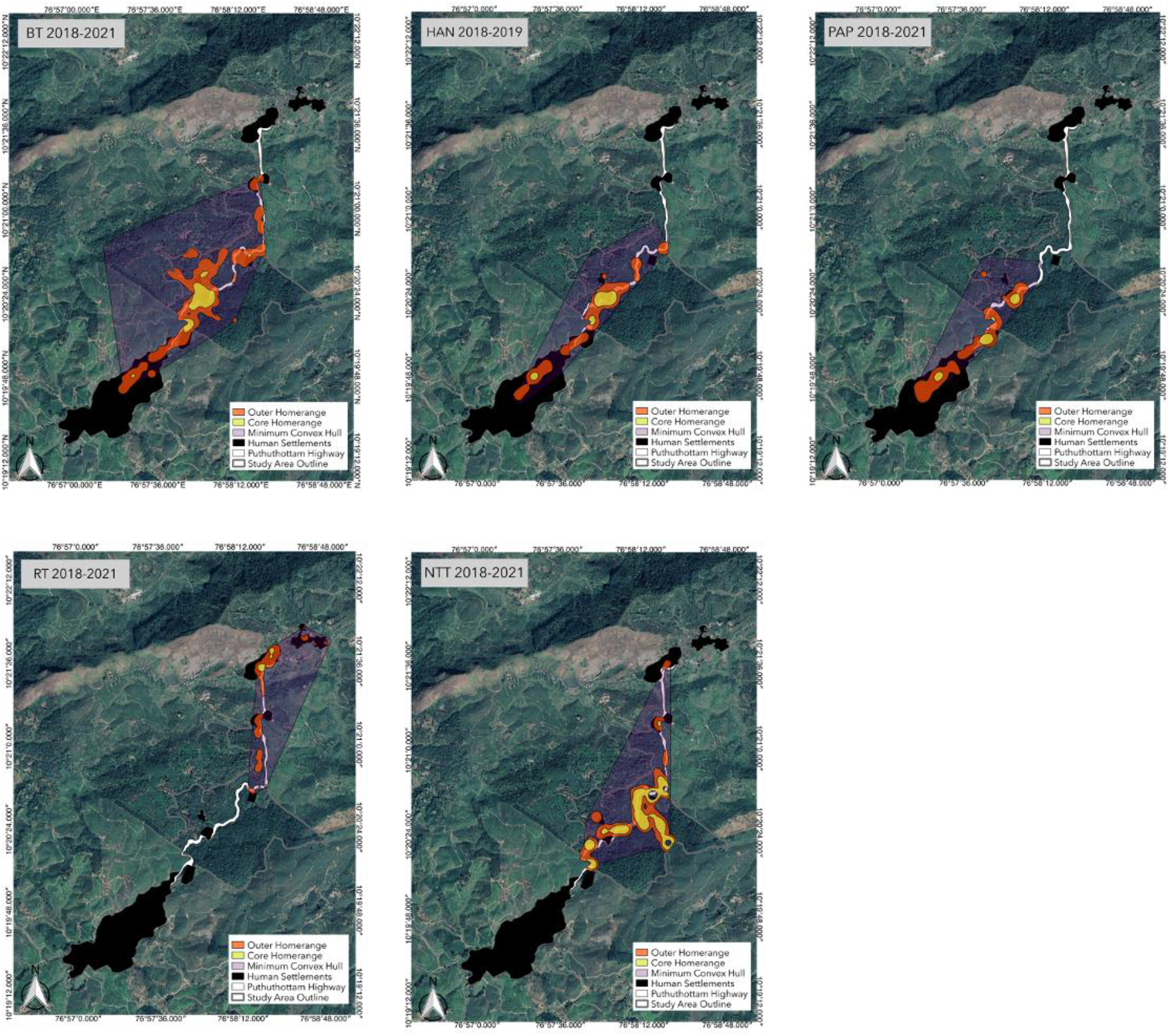
Overall home ranges of all present in Puthuthottam (2018-2021)

**Fig 4.**
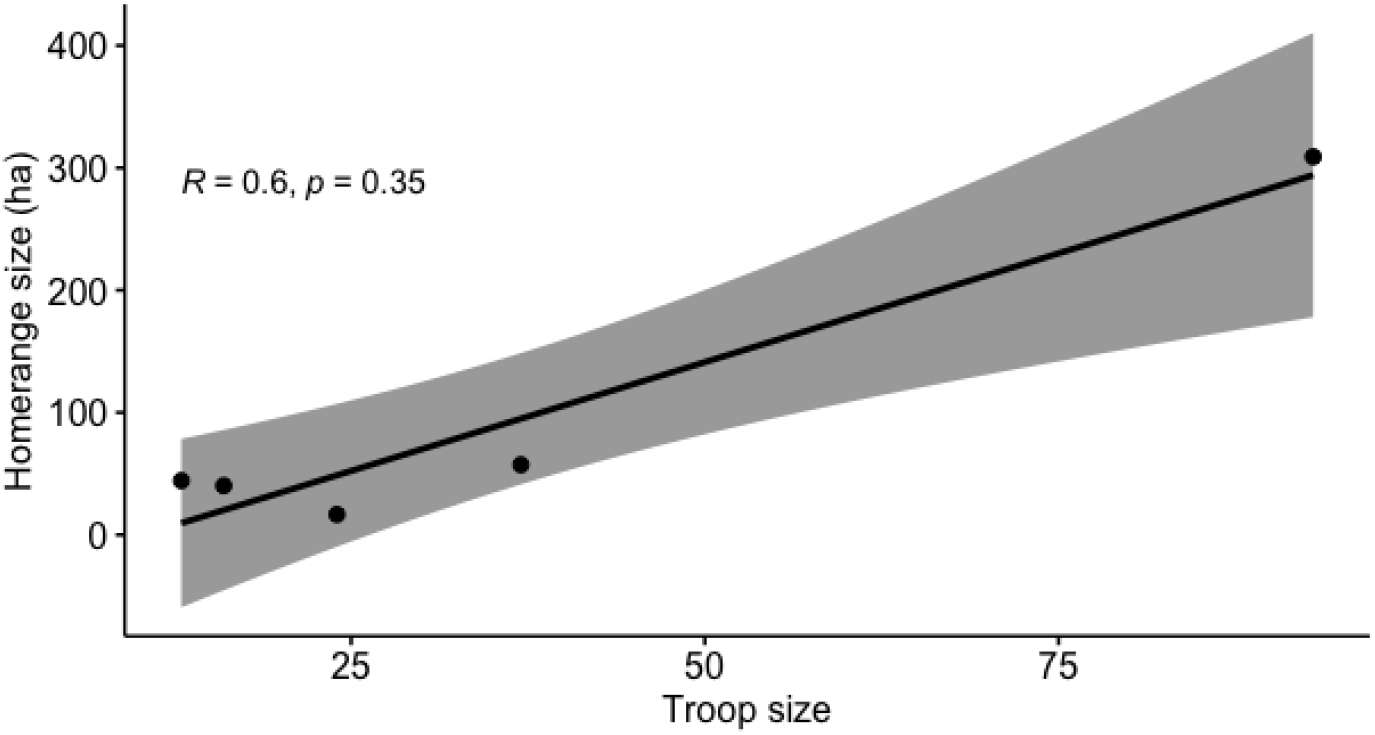
Correlation between total home range area and total troop size for all troops in Puthuthottam

### Home range Overlap

The outer- and core-home ranges of most troops overlapped to a certain degree. There was relatively less overlap between troop core-home ranges than the outer home range. Figures 4 and 5 depict the pairwise overlap of core- and outer-home ranges respectively. Three troops, which were primarily observed in the southern part of the forest fragment, showed the most home range overlap while the two troops near the northern half did not show much overlap.

**Fig 5.**
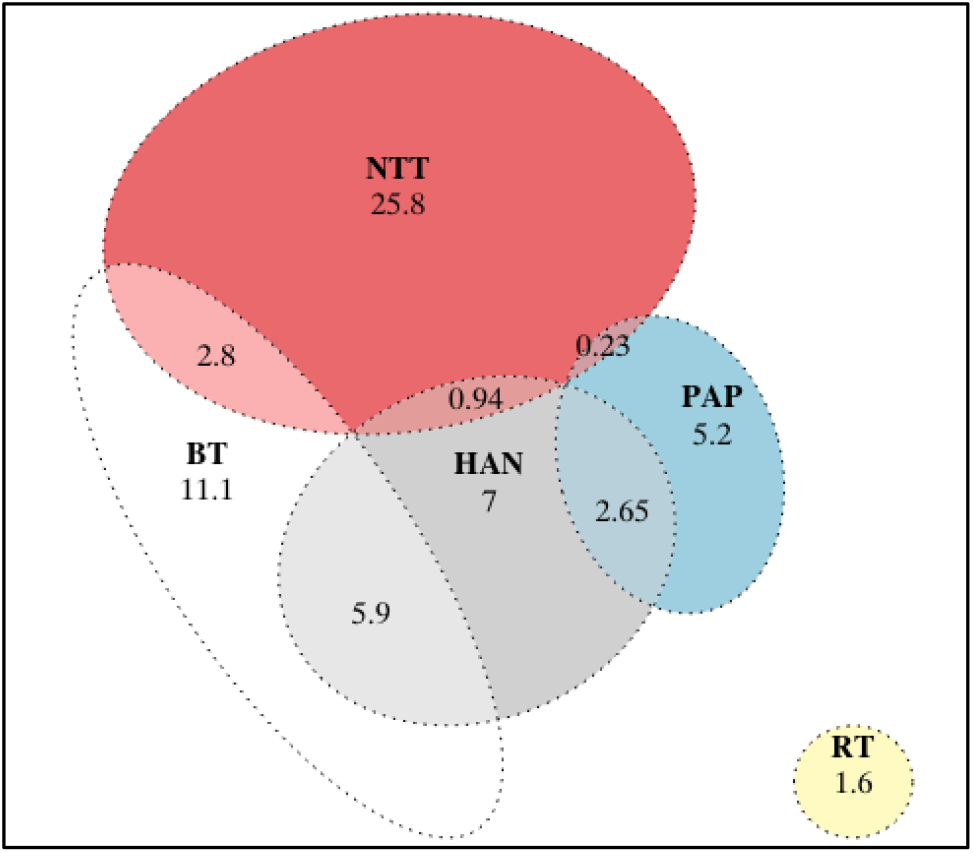
Pairwise overlap of core-home ranges across all troops present in Puthuthottam. Numbers indicate area in hectares

**Fig 6.**
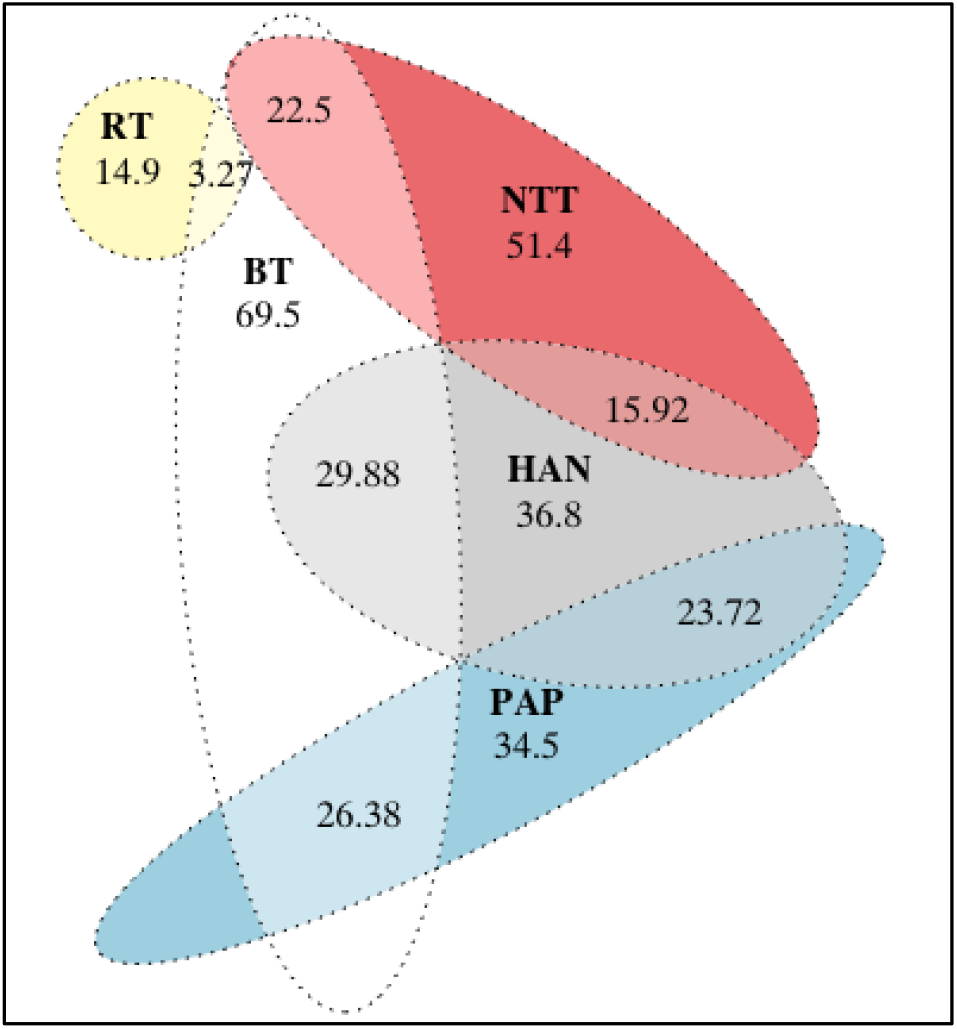
Pairwise overlap of outer-home ranges across all troops present in Puthuthottam. Numbers indicate area in hectares

### Degree of Movement

Over the study period, troops moved an average of 393.3 – 722.3m per daily field session (Table 2). The two troops that joined to form a single troop, NTT, during the first field season moved the most on average per day, however, the largest troop recorded the maximum daily path length at 3.8 km on a single day. Overall, daily path length was not significantly correlated with troop size (R=0.57, p=0.2).

**Table 2.** Observed degree of movement (m) per day for each troop in Puthuthottam 2018-2021

| TROOP ID | AVERAGE OBSERVED MOVEMENT/ DAY (M) | RANGE OF MOVEMENT/ DAY (M) |
| --- | --- | --- |
| BT | 650.5 | 26.5 - 3774.7 |
| HAN | 585.3 | 8.6 - 2467.2 |
| PAP | 393.3 | 6.3 - 1256.4 |
| RT | 468.4 | 4.7 - 2962.9 |
| NTT | 722.3 | 65.1 - 2397.4 |
| IPH | 285.3 | 50.6 - 546 |
| TT | 721.93 | 64 - 2565 |

### Habitat Preference

A non-parametric Wilcoxon test revealed a significant difference between density-dependent randomly generated points and observed macaque locations (W= 24312; p <0.0001; Figure 7). The macaques also used human-dominated habitats such as the road and human settlements disproportionately more than the area available in these habitat types (Figure 8).

**Fig 7.**
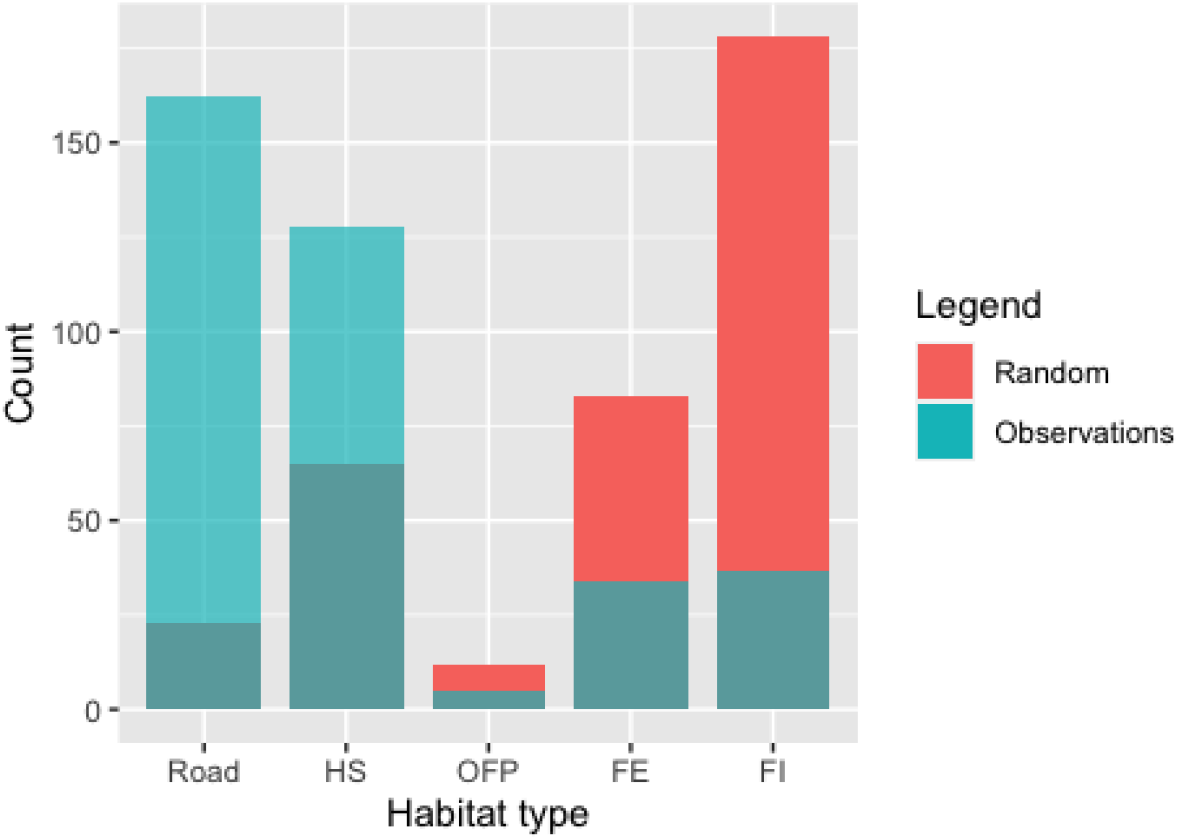
Randomly generated locations versus observed lion-tailed macaque locations across each habitat type. HS= Human Settlement; OFP= Open Forest Patch; FE= Forest Edge; FI= Forest Interior

**Fig 8.**
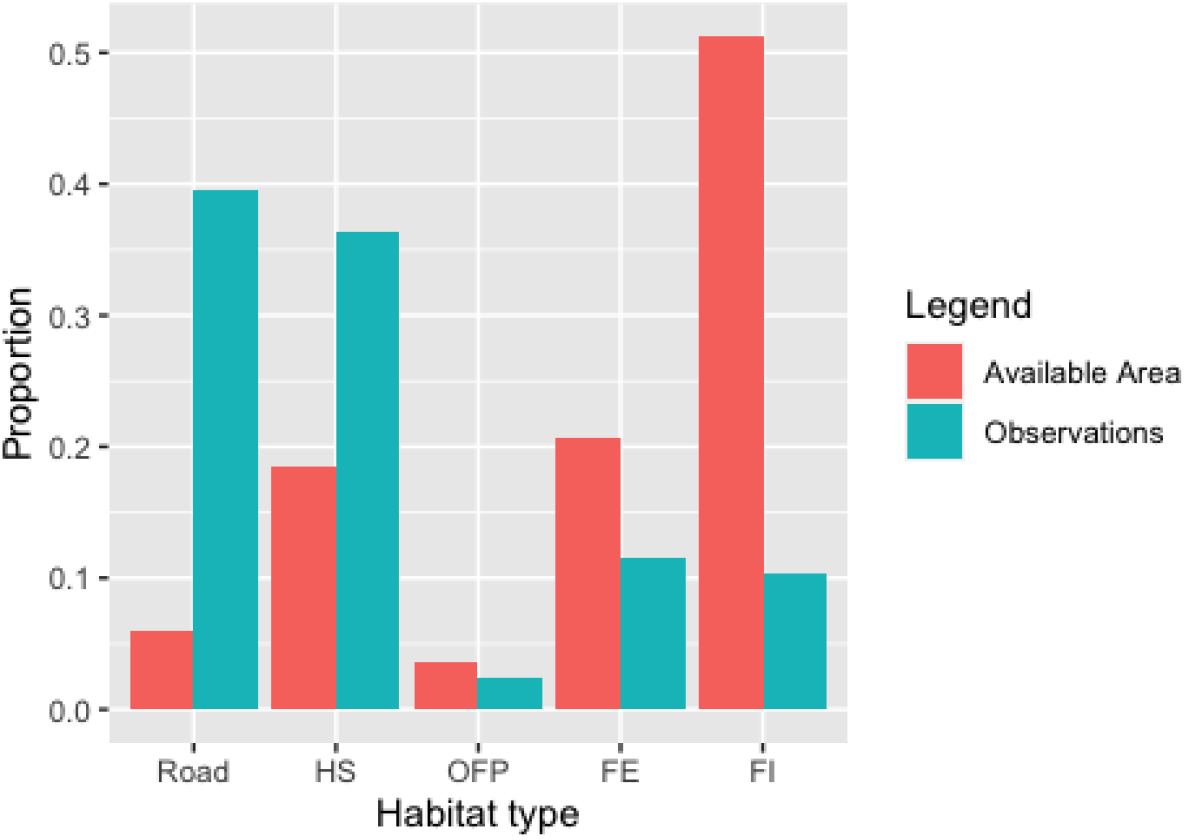
Available area in each habitat type versus observed lion-tailed macaque locations. HS= Human Settlement; OFP= Open Forest Patch; FE= Forest Edge; FI= Forest Interior

## Discussion

Each of the five Puthuthottam troops visited human habitation at varying rates, with the largest troop, BT, visiting most frequently, followed by RT, the newly formed troop in the population. The HAN troop, which split from the BT troop most recently in 2017 also visited human habitations relatively frequently. Interestingly, the two smaller troops, IPH and PAP, which infrequently visited human settlements or the road, both contained a single adult male up until 2016, after which an additional male joined each troop. Both troops were observed to increase habitation visitation after this time, with IPH mainly frequenting the Iyerpadi Garden Hospital and some stretches of road towards the north of the fragment, and PAP moving along the edges of the Valparai town towards the south. The rate at which Puthuthottam monkeys visited houses and buildings was calculated at 0.43/day in 2001 (Singh et al., 2001) while this study indicates an increased habitation visitation rate of 0.57/day. Additionally, our measurement, which required continuous monitoring of the road and settlements, was carried out for a year between 2018-2019; based on our further observations of troops, these rates were noted to increase in the following years between 2019-2021.

All of the five Puthuthottam troops incorporated, into their core-or outer-home range, one or more of the six human settlements situated within and bordering the Puthuthottam forest fragment, listed from North to South as follows: Rottikadai, Iyerpadi Garden Hospital, 10-Acre, Puthuthottam lines, PAP colony and the Valparai town. Energy-rich human-use foods were, naturally, accessible at each of these settlements, however, Rottikadai, Puthuthottam lines and the Valparai town contained large areas where garbage was openly disposed and were presumably the most contested sources for this precious food type. While Rottikadai is a kilometre to the North of the fragment and requires traversing a dangerous highway, tea fields and swamps, the Puthuthottam lines and Valparai are located to the South of the fragment, where we see a corresponding concentration of troop activity. Of the five troops, home ranges of three were located entirely in the southern half of fragment, and the remaining two troops maintained home ranges towards the northern half of the fragment.

Across taxa, home range size is largely determined by diet, body size and corresponding energy requirements (Harestad & Bunnel, 1979; McNab, 1963). Within primate species as well, home range size is dependent on body size and diet, with folivorous and terrestrial species maintaining smaller home ranges than frugivorous and arboreal species (Milton & May, 1976). Additionally, group-size plays an important role in determining home ranges, most primates being group-living, with larger groups maintaining larger home ranges, to fulfil the metabolic requirements of all troop members (Clutton-Brock & Harvey, 1977). The lion-tailed macaque, an arboreal and primarily frugivorous species, has been reported to maintain a home range size ranging between 1.25km (Kumar, 1987) to 5km (Green & Minkowski, 1977) in the wild. In selectively logged forests of Sirsi-Honnavara, the reported home range for lion-tailed macaque groups is a maximum of 3km, with an average daily path length of 500-1500m (Santhosh et al., 2015). The unique habitat composition of the Puthuthottam forest fragment, however, creates a hard boundary beyond which troops are unable to move, due to the presence of impermeable tea plantations and swamps; with the exception of the largest BT troop, which contains c. 96 individuals, the Puthuthottam troops all showed drastically reduced home range sizes ranging between 9.7ha to 98ha. Further, the total home range size did not vary significantly across the five troops, although BT, the largest troop, did maintain the largest home range. Consequently, the daily path length were also observed to be reduced, ranging between 285-722m/day, and were also comparable across troops. The ability for this population to sustain small home ranges, despite requiring much larger areas, is explained by the presence of easily available human-use foods, which allow individuals to acquire greater energy per unit food (Altmann & Muruthi, 1988), thus, resulting in a patterns of altered ranging behaviour also observed in many provisioned species (e.g. Berman et al., 2007; Sengupta et al., 2015; Sinha & Mukhopadhyay, 2013).

Since the Puthuthottam forest fragment restricts macaque movement beyond certain edges, inter-troop encounters are inevitable. In primates, inter-troop encounters are observed to typically be agonistic in nature (Dorothy L Cheney, 1987) as they increase inter-group feeding competition and, thus, directly influence movements of troops (e.g. Spironello, 2001). In this connection, variations in troop size are thought to be of benefit in defending territories, both in terms of food resource and mates (Wrangham, 1980), a theory which supports our observations of a prevailing inter-troop dominance hierarchy in the Puthuthottam population, wherein the smaller troops tend to avoid encounters with the largest BT troop. A similar trend was also observed in Amboseli, where a large troop of vervet monkeys expanded their range over those of smaller troops, restricting them to certain areas (Cheney & Seyfarth, 1987). Previously, the rate of encounters between troops in Puthuthottam has been reported at 0.1/hour, however, this measure has perhaps increased with the increased troop numbers. We, thus, expected this highly competitive environment to have led to scramble competition across the troops in the Puthuthottam, as is often seen with competing primate troops (Isbell, 1991), and evidence from the present study seems to indicate this is indeed the case. Of the three troops that had relatively larger overlaps in home range in the southern part of the study site, two were the smallest troops comprising of 14-16 individuals, allowing them to roam over the same areas without encountering the largest BT troop often. The two northern troops were also able to avoid frequent encounters, especially after one troop migrated entirely out of the forest fragment into neighbouring human settlements, and was able to maintain a core-home range that did not overlap with any other troop.

Finally, while it was evident that human settlements were incorporated in the home ranges of each of the Puthuthottam troops, it was equally important to determine the extent to which these habitat types were being used. Other primate species that have been provisioned often preferentially seek out these resources, thus, gravitating towards human settlements (e.g. Sinha & Mukhopadhyay, 2013). This preference can be an indicator of the degree to which a species is dependent on human-use foods, and its vulnerability to the suite of threats that accompany provisioning. The Puthuthottam troops showed an unfortunate, albeit expected, pattern wherein human-dominated habitats where human-use foods were easily available, such as the Puthuthottam Road and Human Settlement, were used disproportionately more than the available forest habitats, despite these being larger in area. Nevertheless, all troops did use the Forest Interior and Open Forest Patch habitats where they maintained roosting sites throughout the study period. A pertinent point to be made is that despite the troops relying on resources available in human-dominated habitats, the species is still highly dependent on the remaining natural vegetation, without which the population’s survival would be questionable.

Dependence on human-use foods has led to many threats faced by individuals of the Puthuthottam population, some of which are fatal. Singh et al., 2001 reported Puthuthottam troops crossing the main road at 0.7/day, and once again the current measure is perhaps much higher. During the three year study period, five deaths were recorded due to collisions with vehicles on the Puthuthottam Road. Other linear intrusions, such as electric lines passing through the fragment and human settlements, caused two deaths and two minor electrocutions in the population. We also observed numerous injuries to the hands and legs of macaques, most likely from manipulating man-made structures in order to access human-use foods, such as garbage dumpsters, windows and roof tiles. Most importantly, the frequent visits to human settlements has led to a precipitous human-macaque conflict situation, especially in the two settlements, Rottikadai and Puthuthottam lines, where home-raiding occurs most often. A strong presence by the local forest department has averted hunting or retaliatory poisoning cases, however, local community members have continued to build pressure, calling for the capture and translocation of macaques. Historically, such measures have not been successful, leading to the shifting of a problem to a new location, rather than a solution. Furthermore, in all likelihood, macaques from Puthuthottam could culturally transmit home-raiding tendencies and the affinity for human-use foods to neighbouring populations, thus, drastically aggravating the situation. So far, the Puthuthottam population has managed to survive and grow exponentially with the added help of this new food resource, however, our data (see Chapter 2) shows a gradual steadying of the population as resources become increasingly limited. These natural underlying processes would perhaps exert the desired control on the population far better than those offered by further human intervention.

